# Tribbles 3 blunts exercise-induced skeletal muscle adaptation

**DOI:** 10.1101/2025.07.16.665065

**Authors:** Ran Hee Choi, Abigail McConahay, Mackenzie B. Johnson, Morgan Ojemuyiwa, Brooklyn J. Phillips, Ha-Won Jeong, Ho-Jin Koh

## Abstract

Tribbles 3 (TRB3) is a pseudokinase and its expression has been shown to disrupt glucose metabolism through the inhibition of Akt under obese and diabetic conditions. We recently found that overexpression of TRB3 in mouse skeletal muscle decreased skeletal muscle mass and function, leading to muscle atrophy. Here, we examined whether TRB3 affects exercise training-induced skeletal muscle adaptation. We trained muscle-specific TRB3 transgenic (TG) and wild-type (WT) littermates using a voluntary wheel running protocol for 6 weeks and found that TG mice ran significantly less weekly distances than WT littermates. To exclude the possibility that different skeletal muscle adaptations would be produced due to different training intensities, involuntary treadmill exercise (TM) was used as a training regimen. At the 5^th^ week of training, we measured glucose tolerance and found that trained TG mice showed glucose intolerance compared to WT littermates. Furthermore, overexpression of TRB3 significantly suppressed the expression of genes needed for glucose uptake and mitochondrial biogenesis, independent of training status. To further determine the role of TRB3 in exercise-induced adaptation, TRB3 knockout (KO) mice were trained by voluntary and involuntary exercise protocols. KO mice presented improved glucose tolerance compared to WT littermates independent of training status. However, we did not observe significant change in the expression of markers for glucose uptake and mitochondrial biogenesis. Taken together, our results indicate that TRB3 in skeletal muscle blunts the benefits of exercise-induced skeletal muscle adaptation.

## INTRODUCTION

Tribbles 3 (TRB3), a mammalian homolog of Drosophila tribbles, is a pseudokinse, which lacks catalytic function, thus it serves as a molecular scaffold (4). It is responsible for various cellular events, including glucose regulation, endoplasmic reticulum stress response, differentiation of skeletal muscle cells and adipocytes, and cell cycle regulation (12, 20, 32, 33, 37).

For decades, many studies have investigated and confirmed the finding that TRB3 has a negative effect on Akt phosphorylation, which is a major process for glucose and lipid metabolism in liver and skeletal muscle (12, 40). For instance, hepatic TRB3 expression decreases phosphorylation of Akt and impairs glucose tolerance, while TRB3 knockdown improves hepatic glucose regulation (12). In skeletal muscle, muscle-specific TRB3 overexpression worsens high-fat diet-induced obesity and decreases phosphorylation of Akt and glycogen synthase kinase 3 (GSK3), while muscle-specific TRB3 knockout improved glucose homeostasis with high-fat diet (HFD) feeding (21, 40). Our lab recently demonstrated that muscle-specific TRB3 overexpression regulates skeletal muscle mass by altering protein turnover through the regulation of Akt and its downstream molecules, mechanistic target of rapamycin (mTOR) and forkhead box O transcription factors (FOXO) (6, 7). Given the findings that TRB3 could be an important regulator of skeletal muscle mass and integrity, we hypothesized that TRB3 may significantly alter the exercise-induced adaptation to skeletal muscle.

Skeletal muscle, which composes 40-50% of body mass, is a major organ that regulates whole body energy metabolism because it is responsible for ∼80% of postprandial glucose uptake (9). Continuous contractile activity, such as that elicited by exercise, has been associated with improved glucose and oxidative metabolism in skeletal muscle (2, 3, 10). Exercise-induced improvement in glucose metabolism is associated with increased expressions and functions of glucose transporters and metabolic enzymes in skeletal muscle. Previous studies have found that glucose transporter 4 (GLUT4) mRNA expression is increased after exercise (23, 24), but protein expression after exercise varies (13, 25). In addition to increased GLUT4, it is reported that a single bout of exercise increases hexokinase II (HXKII) expression and enzymatic activity in skeletal muscle, which improve glucose metabolism (30, 31). Exercise-induced oxidative capacity and metabolic efficiency are elicited by increased mitochondrial biogenesis (34).

Peroxisome proliferator-activated receptor-gamma coactivator 1α (PGC1α) is highlighted as a key regulator of mitochondrial biogenesis in skeletal muscle in response to endurance exercise (3, 18). PGC1α is a transcription factor that can coordinate the expression of genes related to mitochondrial biogenesis, such as mitochondrial transcription factor A (TFAM), cytochrome oxidase (COX) 1 and 4, and cytochrome c (Cyt c) (39). Previously, muscle-specific PGC1α knockout mice showed impaired muscle function, locomotion, and oxidative capacity, but muscle-specific PGC1α knockout did not prevent exercise-induced COX and Cyt c expression (16, 17).

Increased TRB3 expression has been found in the liver and skeletal muscle from obese and diabetic rodent models, including *ob/ob* and HFD-induced obese mice, which is decreased in response to acute and chronic endurance exercise (26, 28, 29). In addition, reduced TRB3 expression is associated with increased phosphorylation of Akt and improved glucose metabolism (26, 28, 29). One recent study has shown that muscle-specific TRB3 transgenic mice display increased exercise capacity with greater amount of oxidative fibers (1). Although there is some evidence that TRB3 is regulated by exercise, the effect of TRB3 on exercise training-induced skeletal muscle adaptation has not been elucidated.

In the current study, we investigate whether TRB3 in mouse skeletal muscle would disrupt exercise-induced skeletal muscle adaptation, including glucose uptake and mitochondrial biogenesis. Our findings show that muscle-specific TRB3 transgenic (TG) mice run significantly less distances than wild-type (WT) littermates when they are trained by voluntary wheel running. TG mice display significantly suppressed gene expression of GLUT4 and HXKII compared to WT littermates. Furthermore, TRB3 overexpression represses the expression of genes responsible for mitochondrial contents, such as PGC1α, COX4, and Cyt c in both sedentary and trained groups. TRB3 knockout (KO) mice have better glucose tolerance than WT littermates independent of training status, but there is no significant improvement in exercise-induced skeletal muscle adaptation at the molecular levels. Taken together, these results indicate that TRB3 in skeletal muscle blunts the benefits of exercise in glucose uptake and mitochondrial biogenesis.

## MATERIALS AND METHODS

### Animal care

Ten- to fourteen-week-old muscle-specific TRB3 transgenic (TG) and whole-body TRB3 knockout (KO) mice, and their wild-type littermates were used for the current study. All animals were fed standard chow diet (Harlan Teklad Diet, Madision, WI, USA) and accessed water *ad libitum*. Mice were maintained in the animal facility at the University of South Carolina and were housed in an environmentally controlled room on a 12 hr light-dark cycle. All protocols were approved by the Institutional Animal Care and Use Committee of the University of South Carolina.

### Voluntary wheel cage exercise

Mice were randomly assigned to housing in individual cages with or without running wheels (9.5 inch in diameter) for 6 weeks. Mice freely accessed to wheel (Mini Mitter company, Sunriver, OR, USA). Daily activities, including running time and wheel revolutions, were monitored by computer monitoring software Vital View (Starr Life Science Corp., Oakmont, PA, USA). Average daily running distance and 6-week total running distance were calculated from the monitored data using Microsoft excel. Before sacrifice, the wheels were removed from the cage for 48 h to minimize acute exercise effect.

### Involuntary treadmill running protocol

Separated ten- to fourteen-week old TG and WT mice were randomly assigned to treadmill exercise groups. All mice in exercise groups had an acclimation period for 3 days on the treadmill (Columbus instruments treadmill simplex II, Columbus, OH, USA) for 10 min at 8 meters/minute speed. For the 1^st^ week of the training, mice ran for 1 h/day at 12 meters/minutes at 5 degrees incline for 5 days/week. From the 2^nd^ week to last week, running speed was increased up to 16 meters/minutes and remained the same for 6 weeks.

### Glucose tolerance test

At week 5, mice were transported from wheel cages to normal cages and underwent overnight 16 h fasting to determine glucose tolerance. After 16 h fasting, fasting blood glucose was determined. Glucose (2 g/kg body weight) was intraperitoneally injected. Blood glucose was measured by a glucometer (Bayer, Leverkusen, Germany) at 15, 30, 45, 60, 90, and 120 minutes after the injection.

### Muscle glycogen content

Frozen gastrocnemius muscles were pulverized and used to detect glycogen content after exercise training. The content was determined by hydrochloric acid hydrolysis for 2 h at 95 ℃ followed by sodium hydroxide neutralization. Then, the concentration was measured spectrophotometrically using a hexokinase-based assay kit (Sigma, St. Louis, MO).

### Citrate synthase activity

Citrate synthase activity was determined spectrophotometically in gastrocnemius muscle as previously described (36). Gastrocnemius muscles were pulverized and used 5 – 7 mg of muscle to measure the enzyme activity. Homogenates were incubated with substrate and cofactors at room temperature and the working wavelength is 412 nm to detect the reaction product thionitrobenzoic acid.

### Body weight, food intake, blood glucose and body composition

Body weight, food intake, and blood glucose were monitored weekly at the same time.

Body composition was assessed at week 5. Mice were briefly anesthetized by isoflurane inhalation and assessed for percentage fat and lean mass by dual-energy X-ray absorptiometry (DEXA; Lunar PIXmus, Madison, WI).

### Western blot analysis

Frozen gastrocnemius muscles was used to prepare protein lysates for Western blot analysis as previously described (7). Protein lysates (30 µg) were separated on 7 – 15% of SDS-polyacrylamide gels and transferred to PVDF membranes. Primary and secondary antibodies purchased from commercial sources, including GLUT4 (07-1404; Millipore, Billerica, MA, USA), hexokinase II (sc-3521; Santa Cruz Biothechnology Inc., Dallas, TX, USA), PGC1α (ab5441; Abcam, Cambridge, MA, USA), COX4 (4844), Cyt c (4280) from Cell Signaling Technology (Danvers, MA, USA), TRB3 (ST1032; Calbiochem, San Diego, CA, USA), anti-rabbit (NA934; GE healthcare, Pittsburgh, PA, USA) and anti-goat (31400; Thermo Fisher Scientific, Waltham, MA, USA) IgG-horesradish peroxidase conjugated secondary antibodies.

The specific bands were visualized using ECL reagents (NEL105001EA; PerkinElmer, Waltham, MA, USA), and the intensity of bands was quantified using Image Studio Lite (LI-COR Biosciences, Lincoln, NE, USA).

### RNA extraction and RT-PCR

Total RNA was extracted from TA muscles using Trizol reagent (Life Technologies, Carlsbad, CA, USA) according to the manufacturer’s instruction. cDNA was synthesized from 4 mg of total RNA by High Capacity cDNA kit (Life Technologies, Carlsbad, CA, USA). cDNA and primers for real-time polymerase chain reaction (RT-PCR) was carried by SYBR green PCR master mix (Life Technologies, Carlsbad, CA, USA). Primer sequences used to analyze exercise-induced glucose metabolism and oxidative capacity can be found in Table 5.1. All genes were normalized to the level of 18S, a house keeping gene.

### Statistical analysis

Data were expressed as mean ± SEM. Statistical analysis was performed using GraphPad Prism 7 software (La Jolla, CA, USA). Student’s two-tailed unpaired *t* tests were used for comparisons between two groups. One- or two-way ANOVA was conducted to determine significance between more than two group followed by Tukey’s or Bonferroni’s Post hoc analysis. P < 0.05 was considered statistically significant.

## RESULTS

### Muscle-specific TRB3 transgenic mice run shorter distances

We reported in previous study that increased TRB3 in skeletal muscle under stress conditions, including ER stress and HFD, impairs insulin-stimulated Akt phosphorylation and whole-body metabolism, and that TRB3 knockout abolishes the development of HFD-induced insulin resistance (21). Furthermore, our recent study demonstrateed that TRB3 overexpression in mouse skeletal muscle disrupts protein turnover, which in turn worsens food deprivation-induced atrophy (7). Based on these findings, we hypothesized that TRB3 would play a significant role in exercise-induced skeletal muscle adaptation. In order to test our hypothesis, we individually housed muscle-specific TRB3 transgenic (TG) mice and wild-type (WT) littermates in cages in the presence and absence of wheels for 6 weeks. We weekly monitored body weight, food intake, and random blood glucose. No significant difference was observed in any parameters between the groups (Table 2). The percentage of fat and lean mass was determined by DEXA, and no changes were observed between sedentary and voluntary wheel (VW) groups (Table 2). After 6 weeks of training, gastrocnemius muscles were dissected and used for analysis. TRB3 mRNA expression was not significantly altered by VW in either WT or TG mice (Fig. 1A). Intriguingly, we observed that TG mice ran significantly less distances per week than WT littermates (Fig. 1B). These results suggest that muscle-specific overexpression of TRB3 reduces exercise capacity.

**Figure 1.**
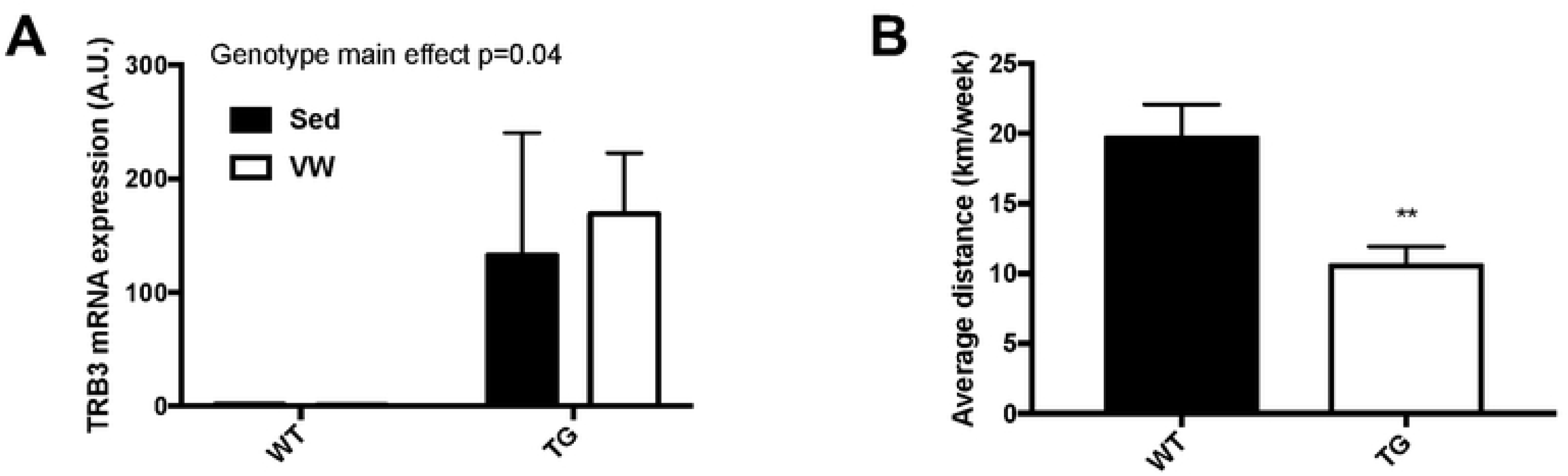
Muscle-specific TRB3 overexpression impairs exercise capacity as measured by 6- week voluntary wheel running. Ten- to fourteen-week old female wild-type (WT) littermates and muscle-specific TRB3 transgenic (TG) mice were separated in sedentary (Sed) and voluntary wheel (VW) running group for 6 weeks. (A) TRB3 expression was determined in gastrocnemius. (B) Wheel activity was monitored by the software and weekly average distance was determined. Data are means ±SEM (n=4/group). ** P<0.01 vs. WT

**Table 1.**
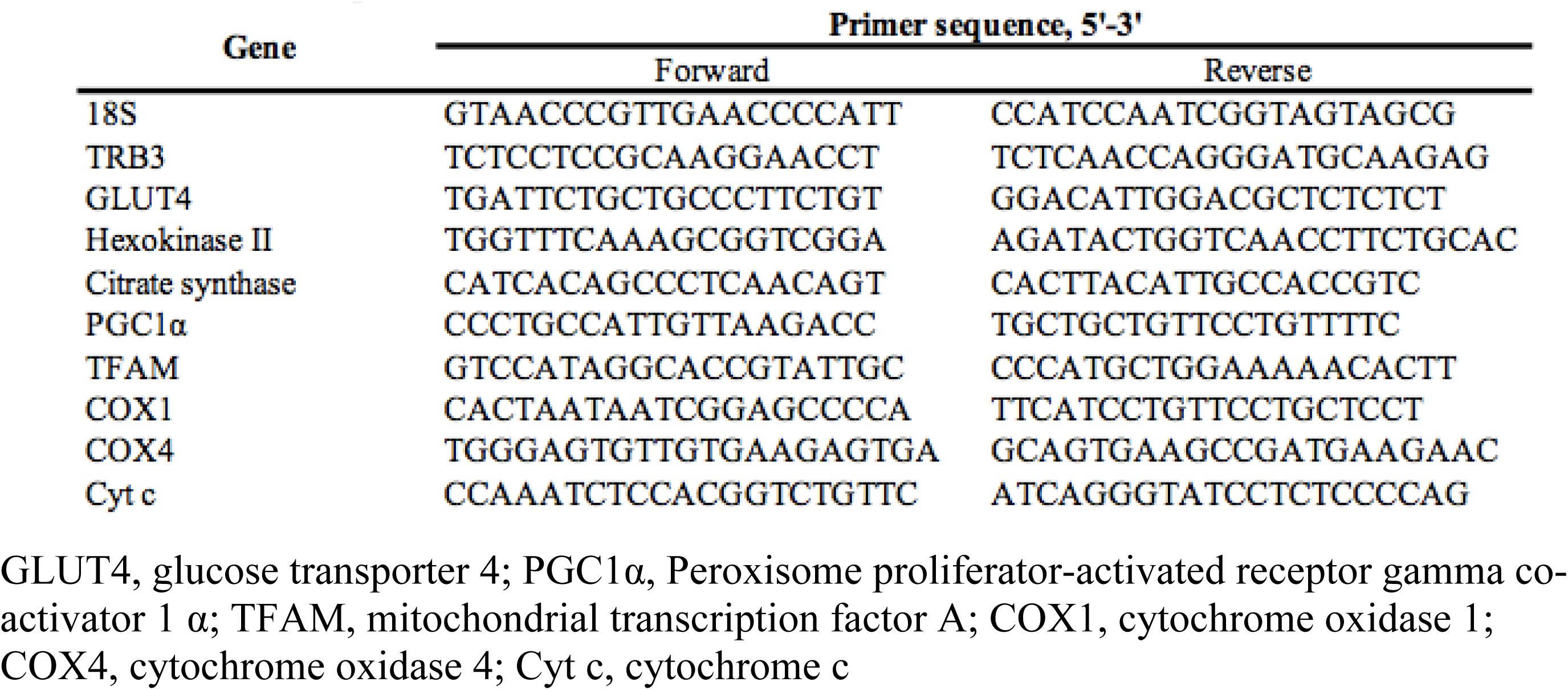
Primer sequences for RT-PCR. GLUT4, glucose transporter 4; PGC1α, Peroxisome proliferator-activated receptor gamma co- activator 1 α; TFAM, mitochondrial transcription factor A; COX1, cytochrome oxidase 1; COX4, cytochrome oxidase 4; Cyt c, cytochrome c

**Table 2.** Characteristics of TG mice in voluntary wheel cage exercise. Ten- to fourteen-week old male female wild-type (WT) littermates and muscle-specific TRB3 transgenic (TG) mice were randomly assigned in two groups; sedentary (Sed) and voluntary wheel (VW) running for 6 weeks. Data are means ±SEM (n=4/group).

| **Characteristics** | **WT** |  | **TG** |  |
| --- | --- | --- | --- | --- |
|  | Sed | Wheel | Sed | Wheel |
| Pre-body weight (g) | 18.6±0.6 | 19.2±0.7 | 18.0±1.1 | 18.1±0.9 |
| Post-body weight (g) | 21.6±0.6 | 22.8±0.8 | 21.4±1.1 | 20.8±0.8 |
| Food intake (g/day) | 4.4±0.2 | 5.1±0.2 | 4.4±0.2 | 4.8±0.3 |
| Blood glucose (mg/dl) | 134.5±9.8 | 129.5±9.5 | 124.8±8.0 | 127.2±10.4 |
| Fat mass (%) | 13.5±0.7 | 15.5±1.0 | 14.3±0.7 | 14.9±0.5 |
| Lean mass (%) | 86.7±0.7 | 84.8±1.1 | 85.9±0.8 | 85.0±0.5 |

### TRB3 overexpression in mouse skeletal muscle does not improve glucose metabolism in response to 6-week treadmill training

Since we observed that TG mice ran significantly less distances compared to WT littermates, we speculated that the difference in training intensities between genotypes could result in different adaptation effects. Thus, we utilized an involuntary exercise protocol, forced treadmill running (TM), to give all mice the same amount of exercise. TG mice and WT littermates were encouraged to run on the treadmill for 6 weeks (details of this procedure can be found in Materials and Methods section). During the 6-week TM training, body weight did not differ between the groups (Table 3). However, we observed that TM groups showed significantly reduced body fat and increased lean mass compared to sedentary groups (Table 3). To investigate whether TRB3 was responsive to forced involuntary exercise, we measured TRB3 mRNA expression. TM running did not alter TRB3 mRNA expression in WT mice, but TG mice showed a greater increase in the mRNA expression in response to the training (Fig. 2A). Next, to determine whether TRB3 overexpression regulates glucose metabolism in response to involuntary exercise, we tested glucose tolerance 5 weeks after the training. In trained WT mice, glucose tolerance was slightly improved, but not by a statistically significance amount (Fig. 2B and C). TG mice in both sedentary and trained groups tended to maintain higher blood glucose during 120-min test periods and showed a higher area under the curve, indicating impaired glucose tolerance (Fig. 2B and C). We further analyzed molecular markers associated with exercise-induced skeletal muscle glucose uptake. Trained WT mice showed no alteration of GLUT4 protein and mRNA expression, while hexokinase II (HXKII) was significantly elevated (Fig. 2D – F). Interestingly, TG mice showed slightly increased GLUT4 and HXKII protein expressions in response to TM training; however, we found that the mRNA expression of both markers was notably suppressed in both sedentary and trained TG mice (Fig. 2D – F). The exercise-induced adaptions in glucose uptake and enzymatic activity has been associated with increased post-exercise glycogen storage (19). Therefore, we measured muscle glycogen contents. No significant difference was detected among groups; however, TG mice tended to show lower amounts of glycogen after training compared to WT littermates (Fig. 2G). These results indicate that TRB3 overexpression in mouse skeletal muscle impairs the benefits of exercise on glucose metabolism by suppressing the expression of genes.

**Figure 2.**
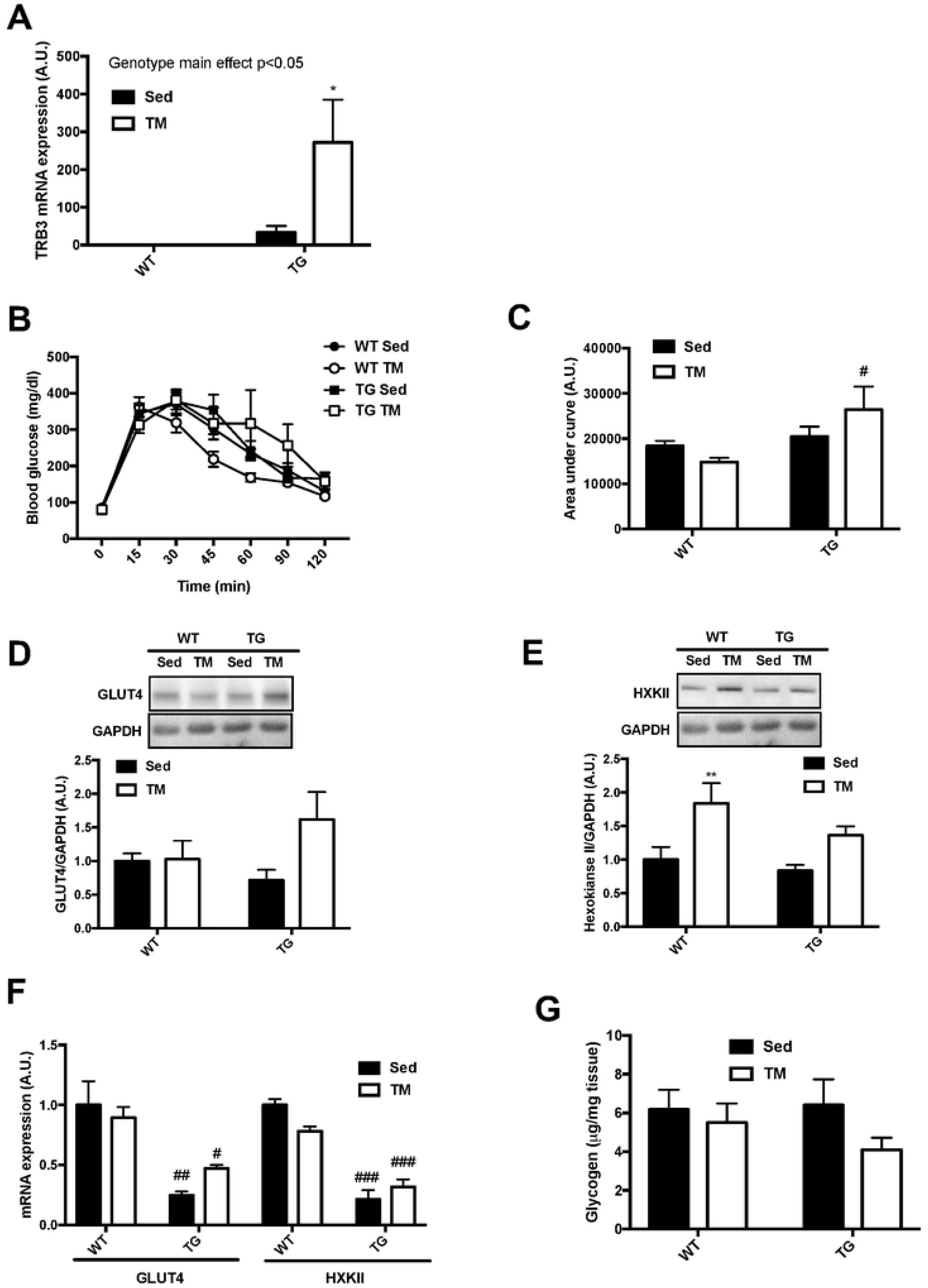
Muscle-specific TRB3 overexpression does not improve glucose uptake in response to 6-week treadmill training. Ten- to fourteen-week old male wild-type (WT) littermates and muscle-specific TRB3 transgenic (TG) mice were randomly assigned in sedentary (Sed) and involuntary treadmill (TM) training groups for 6 weeks. (A) After 6 weeks of training, TRB3 mRNA expression was measured in gastrocnemius muscle. (B-C) At 5^th^ week, glucose tolerance (B) was tested in sedentary (Sed) and treadmill (TM) trained mice and the results were represented as area under the curve (C). (D-E) Protein expressions of glucose transporter 4 (GLUT4; D) and hexokinase II (HXKII; E) were analyzed by Western blot. (F) GLUT4 and HXKII mRNA expression was determined by RT-PCR. (G) Muscle glycogen content were measured in gastrocnemius muscle. Data are means ±SEM (n=4/group). * P<0.05,** P<0.01 vs. Sed; # P <0.05, ## P<0.01, ### P<0.001 vs. WT.

**Table 3.** Characteristics of TG mice in treadmill exercise. Ten- to fourteen-week old male wild-type (WT) littermates and muscle-specific TRB3 transgenic (TG) mice were randomly assigned in two groups; sedentary (Sed) and involuntary treadmill (TM) training for 6 weeks. Data are means ±SEM (n=4/group). * P<0.05 vs. Sed.

| Characteristics | WT |  | TG |  |
| --- | --- | --- | --- | --- |
|  | Sed | TM | Sed | TM |
| Pre-body weight (g) | 25.7±1.0 | 27.33±0.7 | 26.35±0.4 | 27.3±0.7 |
| Post-body weight (g) | 26.7±0.7 | 27.8±0.7 | 26.3±0.6 | 27.9±0.7 |
| Food intake (g/day) | 4.4±0.1 | 4.6±0.2 | 4.3±0.1 | 5.1±0.2 |
| Blood glucose (mg/dl) | 120.5±6.0 | 119.9±9.3 | 118.7±6.4 | 125.7±7.1 |
| Fat mass (%) | 12.7±1.3 | 9.27±0.9* | 14.7±0.7 | 10.9±0.7* |
| Lean mass (%) | 87.3±1.3 | 90.9±0.8* | 85.3±0.9 | 89.2±0.8* |

### Muscle-specific TRB3 overexpression represses exercise-induced mitochondrial biogenesis

Endurance exercise training is known to increase oxidative metabolism in skeletal muscle by increasing the number and size of mitochondria (3, 18). This process is mediated by a transcription coactivator, peroxisome proliferator-activated receptor-gamma coactivator 1α (PGC1α). It has been reported that both acute and chronic exercise can activate PGC1α, resulting in the regulation of other transcription factors related to mitochondrial protein and function (3, 18). Previous research has demonstrated that TRB3 is a mediator of PGC1α-induced insulin resistance in mouse liver (22). Furthermore, muscle-specific PGC1α transgenic mice have shown a paradoxical effect in regard to glucose metabolism and have displayed a two-fold increase in TRB3 expression in their skeletal muscle (5). The previous findings suggest a close relationship between TRB3 and PGC1α in skeletal muscle; however, no research has investigated whether TRB3 expression affects exercise training-induced mitochondrial biogenesis. Therefore, we next tested markers associated with mitochondrial biogenesis and oxidative metabolism. Six-week treadmill training increased the protein expression of mitochondrial biogenesis markers, such as PGC1α, COX4, and Cyt c, in WT mice without altering mRNA expression. (Fig. 3A – D).

**Figure 3.**
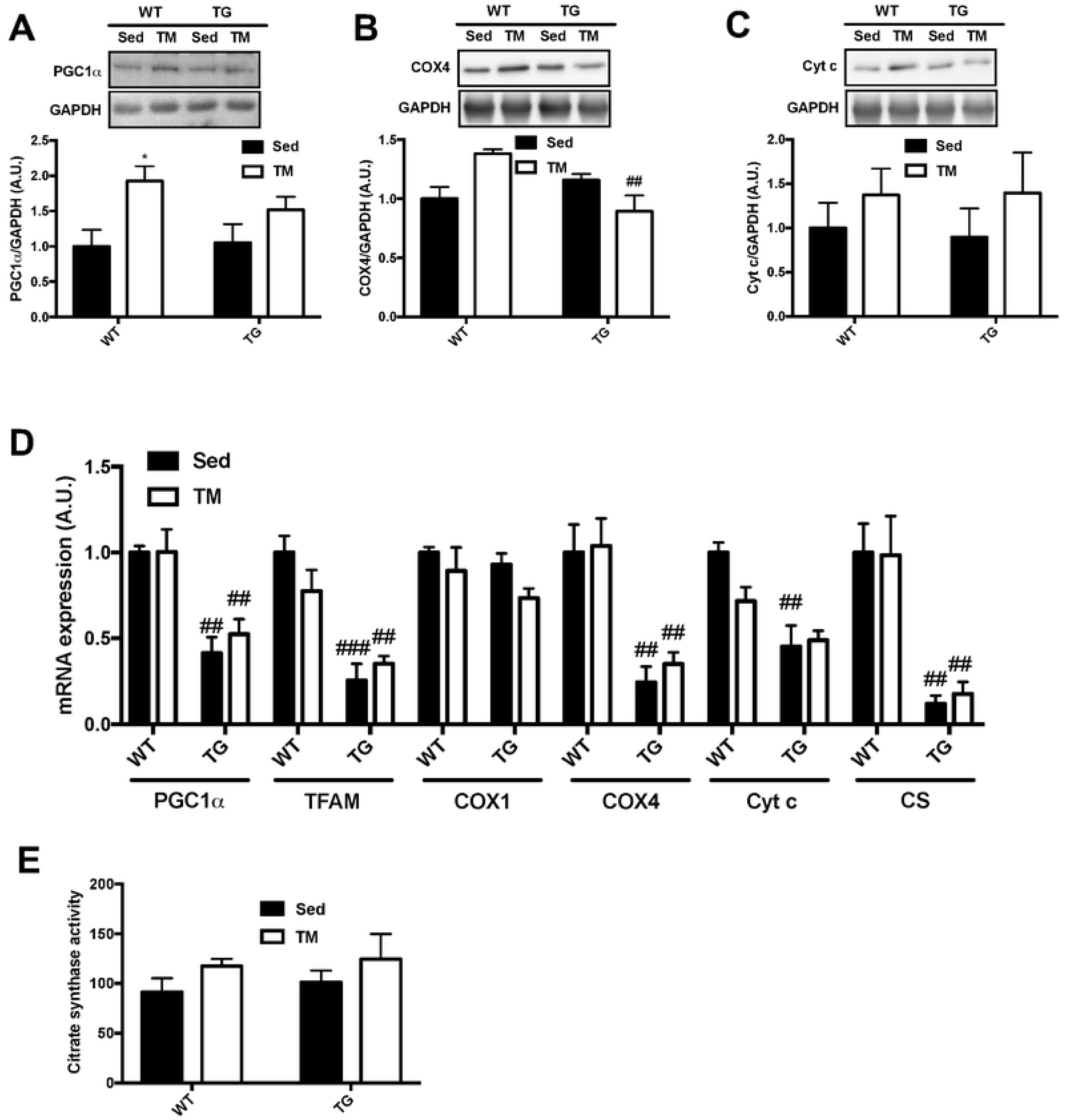
Muscle-specific TRB3 overexpression represses gene expressions of mitochondrial biogenesis markers. Ten- to fourteen-week old male wild-type (WT) littermates and muscle- specific TRB3 transgenic (TG) mice were randomly assigned in sedentary (Sed) and involuntary treadmill (TM) training groups for 6 weeks. (A-C) Protein expression of PGC1α (A), COX4 (B), and Cyt c (C) were determined by Western blot. (D) The expression of genes for mitochondrial contents were measured by RT-PCR. (E) Citrate synthase activity was measured in gastrocnemius to determine oxidative function in response to exercise training. Data are means ±SEM (n=4/group). * P<0.05, ** P<0.01 vs. Sed; # P <0.05, ## P<0.01, ### P<0.001 vs WT.

Furthermore, muscle-specific TRB3 overexpression slightly increased PGC1α and Cyt c protein expression, but significantly reduced COX4 protein expression in trained TG mice compared to trained WT mice (Fig 3. A – C). Intriguingly, TG mice showed dramatically suppressed gene expressions of mitochondrial content markers, such as PGC1α, COX4, and citrate synthase (CS), compared to WT littermates in both sedentary and training conditions (Fig. 3D). Therefore, we tested CS activity to examine whether repressed CS gene expression would affect an enzymatic activity and found that TM training slightly increased CS activity in both groups, which did not reach statistical significance (Fig. 3E). These results indicate that muscle-specific TRB3 overexpression debilitates exercise-induced mitochondrial adaptation by suppressing the expression of genes necessary for mitochondrial biogenesis.

### TRB3 knockout improves glucose tolerance but does not augment the benefits of exercise in glucose metabolism

In the present study, we found that TRB3 overexpression in mouse skeletal muscle blunted exercise training-induced skeletal muscle adaptation by repressing gene expression. Therefore, we considered that a deletion of TRB3 in skeletal muscle would enhance exercise-induced skeletal muscle adaptation. To examine the hypothesis, we next recruited TRB3 knockout (KO) mice and WT littermates for exercise training with voluntary wheel (VW) running (details found in the Materials and Methods section). During 6-week VW training, there were no significant changes in body weight, food intake, blood glucose, or body composition among the groups (Table 4). KO mice ran slightly more than WT littermates, but the increase did not reach to a statistical difference (Table 4). To examine the effect of TRB3 knockout on exercise-induced improvement in glucose metabolism, we measured glucose tolerance after overnight fasting at the 5^th^ week of the training. Exercise training did not affect glucose tolerance in WT and KO mice during the120-min test period (Fig. 4A). Interestingly, trained KO mice showed significantly lower blood glucose than trained WT mice at the 60-min time (Fig. 4A).

**Figure 4.**
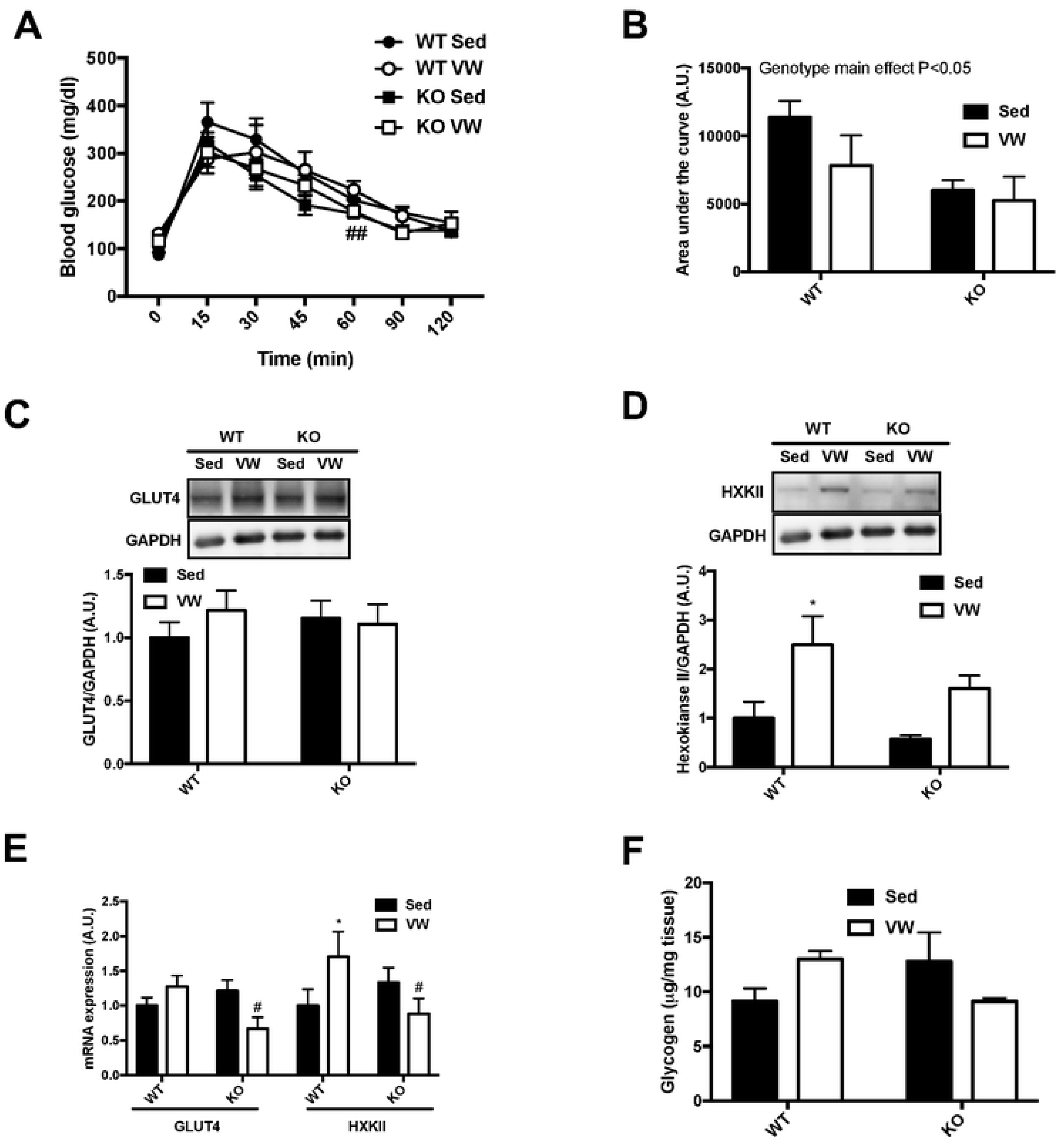
TRB3 knockout improves glucose tolerance but does not augment the benefits of exercise in glucose uptake. Ten- to fourteen-week old female wild-type (WT) littermates and TRB3 knockout (KO) mice were divided into sedentary (Sed) and voluntary wheel (VW) training groups for 6 weeks. (A-B) At 5^th^ week of training, glucose tolerance (A) was measured to determine glucose metabolism and the results were represented as area under the curve (B). (C-D) Protein expressions of glucose transporter 4 (GLUT4; C) and hexokinase II (HXKII; D) were analyzed by Western blot. (E) GLUT4 and HXKII mRNA expression was determined by RT-PCR. (F) Muscle glycogen content were measured in gastrocnemius muscle. Data are means ±SEM (n=4-5/group). * P<0.05 vs. Sed; # P <0.05, ## P<0.01 vs. WT.

**Table 4.** Characteristics of KO mice in voluntary wheel exercise. Ten- to fourteen-week old female wild-type (WT) littermates and TRB3 knockout (KO) mice were randomly assigned in two groups; sedentary (Sed) and voluntary wheel (VW) training for 6 weeks. Data are means ±SEM (n=4-5/group).

| **Characteristics** | **WT** |  | **KO** |  |
| --- | --- | --- | --- | --- |
|  | Sed | Wheel | Sed | Wheel |
| Pre-body weight (g) | 16.9±0.8 | 18.1±0.5 | 17.7±0.3 | 16.6±0.7 |
| Post-body weight (g) | 19.5±0.6 | 20.6±0.5 | 20.2±0.4 | 19.4±0.6 |
| Food intake (g/day) | 4.5±0.1 | 4.8±0.2 | 4.6±0.1 | 4.5±0.1 |
| Blood glucose (mg/dl) | 134±7.3 | 139.3±9.0 | 141.7±8.1 | 138.7±9.1 |
| Fat mass (%) | 13.4±1.1 | 12.9±0.8 | 15.1±0.9 | 12.3±0.5 |
| Lean mass (%) | 86.5±1.1 | 87.0±0.7 | 84.9±0.9 | 87.7±0.5 |
| Average distance (km/week) |  | 11.2±2.4 |  | 14.2±2.3 |

**Table 5.** Characteristics of KO mice in treadmill exercise. Ten- to fourteen-week old male wild-type (WT) littermates and TRB3 knockout (KO) mice were randomly assigned in two groups; sedentary (Sed) and involuntary treadmill (TM) training for 6 weeks. Data are means ±SEM (n=4/group). # P<0.05 vs. WT

| Characteristics | WT |  | KO |  |
| --- | --- | --- | --- | --- |
|  | Sed | TM | Sed | TM |
| Pre-body weight (g) | 27.5±0.6 | 27.3±0.8 | 27.8±1.6 | 25.2±0.7 |
| Post-body weight (g) | 27.8±0.3 | 29.9±0.9 | 28.0±1.7 | 28.6±1.1 |
| Food intake (g/day) | 4.4±0.2 | 4.9±0.2 | 4.5±0.2 | 5.1±0.3 |
| Blood glucose (mg/dl) | 124.6±9.0 | 113.0±5.6 | 121.5±7.3 | 120.0±4.1 |
| Fat mass (%) | 12.1±0.7 | 15.2±1.6 | 13.6±1.6 | 9.8±1.0<sup>#</sup> |
| Lean mass (%) | 87.9±0.7 | 85.1±1.7 | 86.4±1.6 | 89.9±1.1 |

These results were clearly reflected in the area under the curve: TRB3 knockout tended to improve glucose tolerance in sedentary animals and after training (Fig. 4B). To investigate the possible mechanism for improved glucose tolerance in KO mice, we measured GLUT4 and HXKII expressions. In WT littermates, there was no change in GLUT4 protein and mRNA expression, but HXKII was greatly increased in response to VW exercise (Fig 4C – E). VW running did not alter GLUT4 expression in KO mice but did slightly increase HXKII protein expression (Fig 4C and D). Furthermore, KO mice displayed significantly reduced mRNA expression of GLUT4 and HXKII in response to VW training (Fig. 4E). Muscle glycogen content was slightly increased in WT mice after training (Fig. 4F). KO mice tended to show more glycogen content in sedentary status, but exercise training produced no other effects (Fig. 4F). These results suggest that TRB3 knockout improves glucose tolerance in sedentary and trained conditions, but that the improved glucose tolerance is not due to changes in GLUT4 or HXKII.

### TRB3 knockout does not facilitate mitochondrial adaptation in response to VW exercise training

We observed that TRB3 overexpression in mouse skeletal muscle significantly abrogated the expression of genes responsible for mitochondrial biogenesis (Fig. 3D). To determine whether TRB3 knockout accelerates the regulation of exercise-induced mitochondrial adaptation, we examined mitochondrial markers at the protein and mRNA levels. Six-week VW training increased mitochondrial markers, including PGC1α, COX4, and Cyt c, in both trained WT and KO mice compared to sedentary groups (Fig. 5A – C). Consistent with protein expression, trained WT mice showed greatly increased mRNA expression of mitochondrial biogenesis markers, such as PGC1α, COX1 and 4, and CS (Fig. 5D). Surprisingly, TRB3 knockout mice did not show any significant change in the expression of genes needed for mitochondrial biogenesis in response to VW (Fig. 5D). To further examine whether TRB3 knockout affects mitochondrial enzymatic activity, we measured CS activity and found a notable increase in the activity after VW running; however, TRB3 knockout did not further affect its activity (Fig. 5E). These results indicate that TRB3 knockout does not further promote mitochondrial biogenesis markers after VW exercise training.

**Figure 5.**
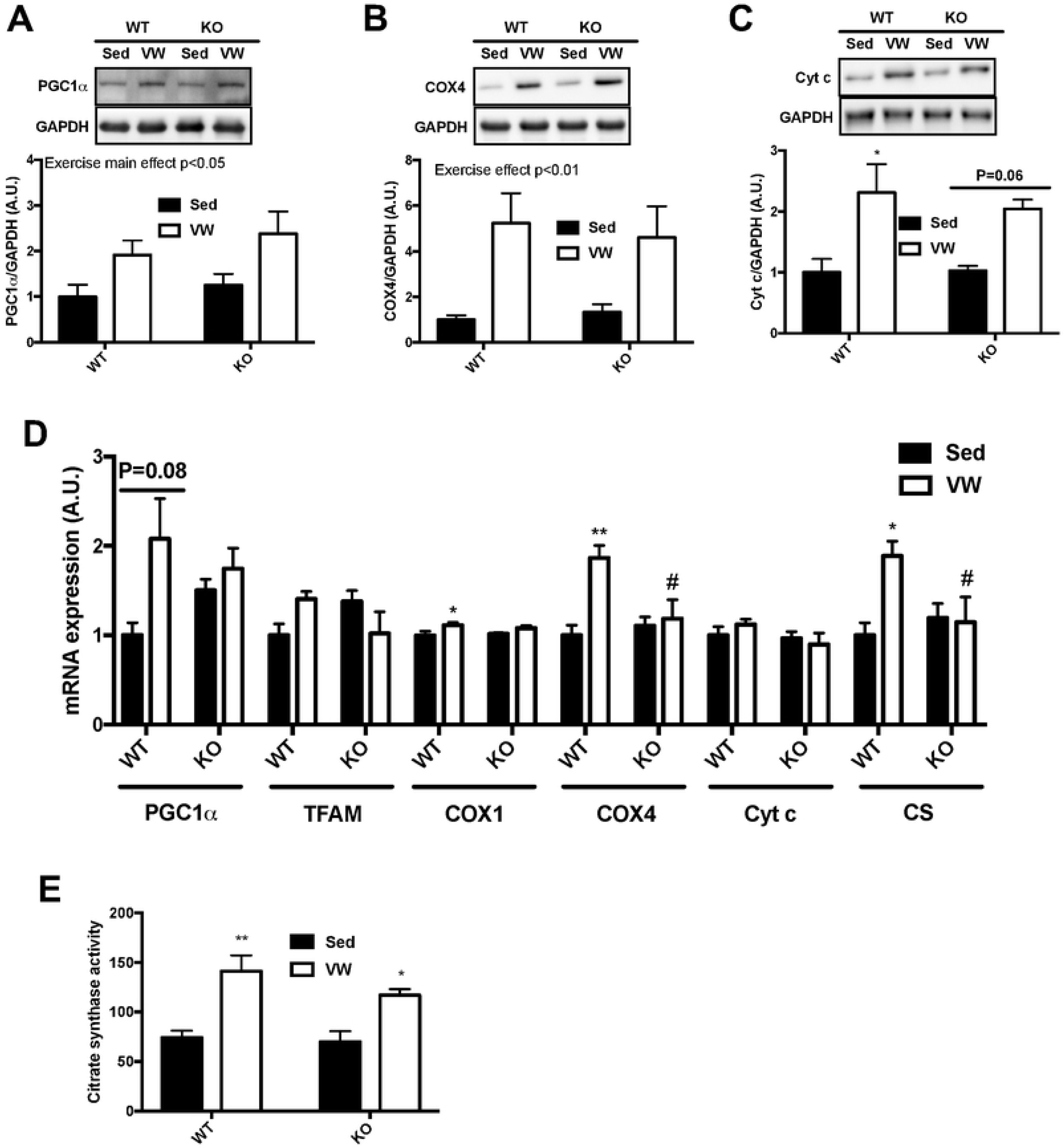
TRB3 knockout dose not facilitate mitochondrial adaptation in response to VW training. Ten- to fourteen-week old female wild-type (WT) littermates and TRB3 knockout (KO) mice were divided into sedentary (Sed) and voluntary wheel (VW) training groups for 6 weeks. After 6 weeks of training, gastrocnemius muscles were analyzed. (A-C) Protein expression of PGC1α (A), COX4 (B), and Cyt c (C) were determined by Western blot. (D) The expression of genes for mitochondrial contents were measured by RT-PCR. (E) Citrate synthase activity was measured to determine oxidative function in response to exercise training. Data are means ±SEM (n=4-5/group). * P<0.05, ** P<0.01 vs. Sed; # P <0.05, ## P<0.01, ### P<0.001 vs WT.

### TRB3 knockout does not benefit exercise training-induced metabolic adaptations in response to involuntary running

In the present study, we have found that TRB3 knockout did not further assist exercise-induced skeletal muscle adaptation even though KO mice were likely to show a better glucose tolerance than WT mice (Fig. 4A and B). To exclude the possibility of the mice receiving different intensities of exercise due to the difference in running distance during voluntary exercise training, we repeated the experiment using an involuntary exercise protocol, forced treadmill running (see Materials and Methods section for details). There were no differences in body weight, food intake, or blood glucose among the groups during the 6-week training period (Table 5). However, we did observe a significant reduction in fat mass in KO mice after TM training compared to WT mice (Table 5). Next, the mice underwent a glucose tolerance test at the 5^th^ week of the training. There was no noticeable difference in glucose tolerance among the groups (Fig. 6A). In KO mice, both sedentary and trained mice showed a lower area under the curve compared to WT mice, suggesting improved glucose tolerance (Fig. 6B). We also analyzed GLUT4 and HXKII to determine possible mechanisms for improved glucose tolerance. GLUT4 and HXKII were increased in response to the involuntary running protocol independent of genotypes (Fig. 6D – E). Furthermore, TM running slightly depleted muscle glycogen in both genotypes, but we did not find any statistical difference (Fig. 6F). Mitochondrial contents at the protein and mRNA levels showed a tendency to respond to TM running, but there were no remarkable differences between genotypes (Fig. 7A – D). In CS activity, TRB3 knockout increased CS activity in response to TM exercise while WT littermates did not show any response to the training (Fig. 7E). Taken together, these data suggest that TRB3 knockout improves glucose tolerance regardless of training status or types of training, but TRB3 knockout does not benefit exercise-induced skeletal muscle adaptation.

**Figure 6.**
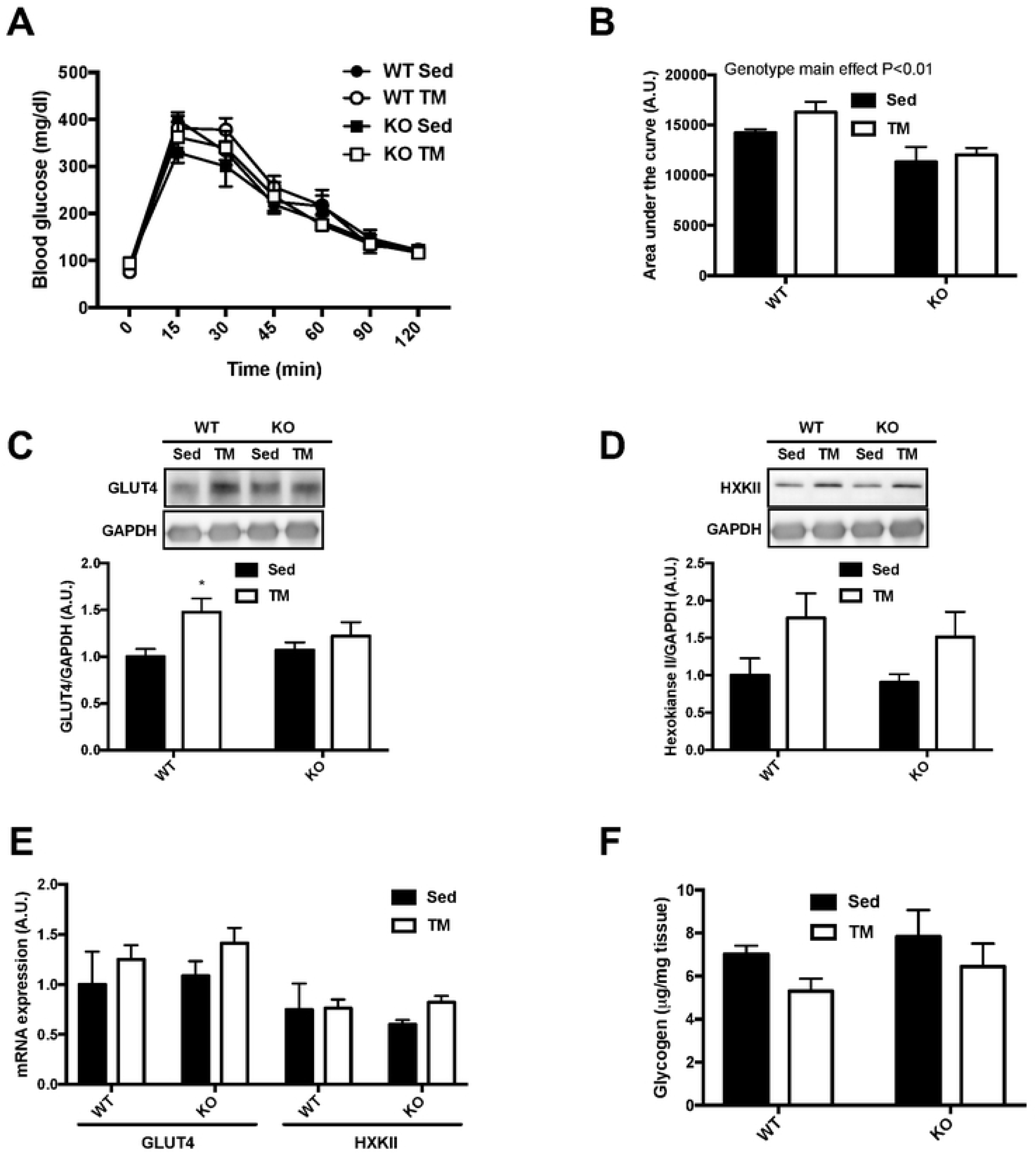
TRB3 knockout does not further improve molecular markers for glucose uptake in response to involuntary exercise. Ten- to fourteen-week old male wild-type (WT) littermates and TRB3 knockout (KO) mice were divided into sedentary (Sed) and involuntary treadmill (TM) training groups for 6 weeks. (A-B) At 5^th^ week of TM training, glucose tolerance (A) was measured to determine glucose metabolism and the results were represented as area under the curve (B). (C-D) Protein expressions of glucose transporter 4 (GLUT4; C) and hexokinase II (HXKII; D) were analyzed by Western blot. (E) GLUT4 and HXKII mRNA expression was determined by RT-PCR. (F) Muscle glycogen content were measured in gastrocnemius muscle. Data are means ±SEM (n=4/group). * P<0.05 vs. Sed.

**Figure 7.**
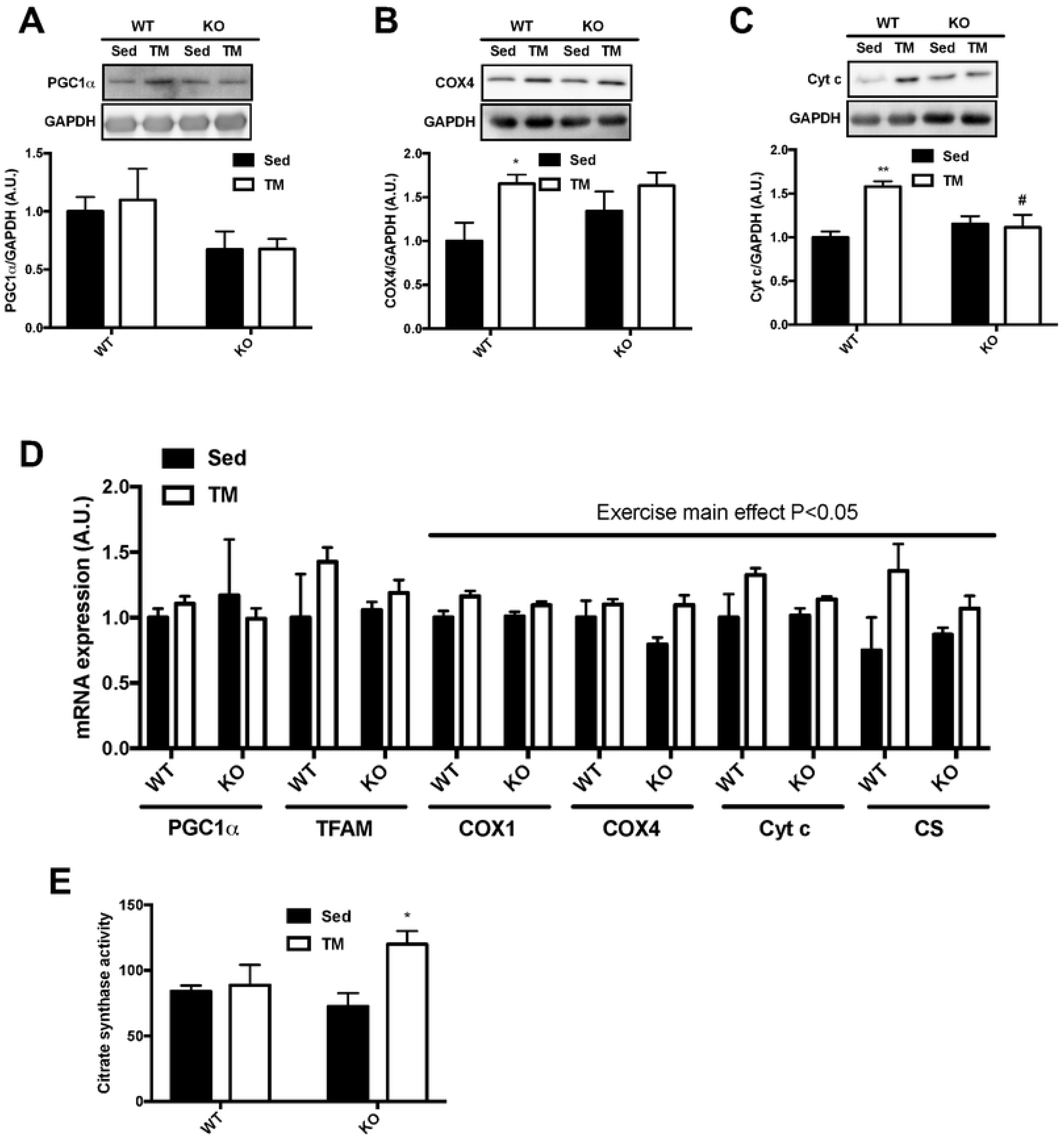
TRB3 knockout does not promote mitochondrial biogenesis in response to involuntary treadmill exercise. Ten- to fourteen-week old female wild-type (WT) littermates and TRB3 knockout (KO) mice were divided into sedentary (Sed) and forced treadmill (TM) training groups for 6 weeks. After 6 weeks of training, gastrocnemius muscles were analyzed. (A-C) Protein expression of PGC1α (A), COX4 (B), and Cyt c (C) were determined by Western blot. (D) The expression of genes for mitochondrial contents were measured by RT-PCR. (E) Citrate synthase activity was measured to determine oxidative function in response to exercise training. Data are means ±SEM (n=4-5/group). * P<0.05, ** P<0.01 vs. Sed; # P <0.05 vs WT.

## DISCUSSION

TRB3 is known to inhibit Akt phosphorylation, and increased TRB3 expression is associated with metabolic dysfunction in multiple tissues, including the liver and skeletal muscle (22, 41). In addition, reduced TRB3 expression in obese and diabetic mouse models has been observed in response to acute and chronic exercise, resulting in amelioration of metabolic stress (26, 28, 29). Although research has demonstrated that TRB3 is strongly associated with metabolic dysregulation in the liver under obese and diabetic conditions, the role of TRB3 in skeletal muscle has not been clearly understood. Our previous studies demonstrated that TRB3 plays important roles in the development of HFD-induced insulin resistance and protein turnover in skeletal muscle at the basal level and under food deprivation-induced atrophy (6, 7, 21). To further determine the role of TRB3 in skeletal muscle, we examined the effect of TRB3 on exercise training-induced skeletal muscle adaptation with regards to glucose uptake and mitochondrial biogenesis. Our findings indicate that muscle-specific TRB3 overexpression impairs glucose tolerance and also suppresses the expression of genes responsible for glucose uptake and mitochondrial biogenesis regardless of exercise training. By contrast with the detrimental effects of TRB3 overexpression, TRB3 knockout improves glucose tolerance in both sedentary and trained groups; however, there were no further improvements in glucose uptake and mitochondrial biogenesis between genotypes in response to exercise training. These findings suggest a possible role for TRB3 in exercise-induced skeletal muscle adaptation: the regulation of whole-body metabolism and oxidative capacity.

In the current study, we observed that voluntary wheel running did not alter TRB3 expression in either WT or TG mice, and involuntary treadmill training greatly increased TRB3 expression in TG mice only (Fig. 1A and 2A). Previously, An et al. reported that 4-week voluntary exercise increases TRB3 mRNA and protein expression in C57BL/6 WT mice (1).

However, we did not observe an increase in TRB3 expression in WT littermates in response to both exercise training protocols. This inconsistency may be due to different types of training procedures, intensity, and duration. Our mice ran about 20 km/week for 6 weeks compared to about 46 km/week for 4 weeks based on An et al.’s report. This difference in running distance raises the possibility of different responses. Additional study may be required using similar intensity, duration, and training protocols to produce comparable results.

Exercise training promotes glucose uptake in skeletal muscle by increasing GLUT4 expression and HXKII enzymatic activity (8, 11, 35). Several previous studies have reported that GLUT4 protein and mRNA expression can vary due to different experiment settings and exercise intensities (13, 24, 25). In the current study, we found that GLUT4 protein expression was not altered by exercise training in either genotype (Fig. 2D), but TRB3 overexpression significantly decreased gene expression of GLUT4 in sedentary and trained groups (Fig. 2F). TRB3 overexpression also repressed HXKII mRNA expression (Fig. 2F). HXKII is a rate-limiting enzyme in glycolysis known to be responsible for exercise-induced glucose uptake (30, 31).

Furthermore, impaired glucose uptake and exercise capacity have been observed in HXKII knockout mice in response to exercise training (14, 15). Given the importance of GLUT4 and HXKII in glucose uptake, suppressed GLUT4 and HXKII gene expression in mouse skeletal muscle may explain the impaired glucose tolerance in TG mice (Fig. 2B and C). Moreover, the decreased gene expression in TG mice may impair exercise capacity and disrupt energy substrate utilization (Fig. 1B). The mechanism(s) by which TRB3 regulates the expressions of genes responsible for metabolism remains to be answered.

Endurance exercise training also stimulates mitochondrial biogenesis. PGC1α is considered to be a main regulator of exercise-induced mitochondrial biogenesis (3). Muscle-specific overexpression of PGC1α increases slow-type fibers and improves exercise capacity (27, 38), while muscle-specific PGC1α knockout mice show decreased daily locomotion and attenuated exercise-induced mitochondrial biogenesis (17). We observed that TG mice displayed significantly lower PGC1α mRNA expression with no further increment after exercise training, and that they also show remarkably reduced expression of genes necessary for mitochondrial biogenesis independent of training status (Fig. 3D). In addition, it has been demonstrated that PGC1α overexpression in liver and skeletal muscle impairs glucose homeostasis by elevating hepatic TRB3 expression (5, 22). Although the role of TRB3 in regulating PGC1α expression has not been elucidated, we believe our results offer at least the possibility that overexpression of TRB3 impairs PGC1α expression and disrupts the regulation of exercise-induced mitochondrial biogenesis.

In the present study, TRB3 knockout mice seem to demonstrate improved glucose tolerance compared to WT littermates, independent of training status (Fig. 5B and 6B).

Previously, our lab has shown that TRB3 knockout prevents HFD-induced development of insulin resistance (21). This would be the first observation of TRB3 knockout possibly benefiting whole-body glucose tolerance on a chow diet, regardless of exercise training. However, we did not detect any further changes in protein or mRNA expressions of GLUT4 or HXKII (Fig. 4C – E, 5C – E). These results suggest that improved glucose tolerance in KO mice is possibly due to other mechanisms than GLUT4 and HXKII, which can regulate glucose metabolism.

Taken together, our data suggest that muscle-specific overexpression of TRB3 impairs exercise capacity and glucose tolerance in response to exercise training by regulating the expression of genes responsible for glucose uptake and mitochondrial biogenesis. We have also demonstrated that the lack of TRB3 improves glucose tolerance, independent of training status. The current study elucidates the possibility that TRB3 expression could affect glucose metabolism and mitochondrial biogenesis in response to exercise training. Additional works are necessary to find the molecular mechanism(s) by which TRB3 regulates transcriptional activity of the genes responsible for glucose uptake and mitochondrial biogenesis.

## ACKNOWLEDGEMENTS

We thank M. Montminy (Salk Institute) for providing the muscle-specific TRB3 transgenic mice, and Regeneron for TRB3 knockout mice. This work was supposed by NIH grants to H.J.K. (R16GM149457, P50CA236733, U54CA163069, and R03AR066825).

## AUTHOR CONTRIBUTIONS

R.H.C. designed the research, performed the experiments, analyzed the data, and wrote the manuscript. A.M., M.B.J., and H.W.J. performed the experiments and edited the manuscript.

M.O. and B.J.P. edited the manuscript. H.J.K. designed the research, analyzed the data, and wrote the manuscript.

